# Transcriptional architecture and Pol II regulation at promoters, enhancers, and enhancer clusters in *Canis lupus familiaris*

**DOI:** 10.1101/2025.03.20.644285

**Authors:** Samu V. Himanen, Adelina Rabenius, Serhat Aktay, Joël Tekoniemi, Anniina Vihervaara

## Abstract

Domestic dog exhibits remarkable phenotypic diversity and provides versatile models for genomics, evolution, and complex traits. DNA sequences and stable RNAs have revealed regulatory regions in the dog genome. However, transcriptional activity, regulatory architecture, and control of RNA Polymerase II (Pol II) across genes and enhancers remain uncharacterized. Here, we track transcription at nucleotide-resolution, measure RNA expression and stability, and analyse mechanisms of Pol II regulation in golden retriever macrophages. We report the precise architectures of promoters, enhancers, and enhancer clusters, and quantify Pol II progression from the initiation, through the pause-region, into elongation and termination. Triggering transcriptional change by heat stress reveals instant reprogramming of genes *via* promoter-proximal pause-regulation and enhancers *via* initiation. Enhancers within a cluster mount a unified response. This study identifies functional genomic regions *de novo*, characterizes transcriptional architectures of genes, enhancers, and enhancer clusters, quantifies RNA synthesis and stability, and reveals mechanisms of transcription in *Canis lupus familiaris*.

## Introduction

Since the domestication of wolf (*Canis lupus lupus*), estimated 30.000 years ago, domestic dog (*Canis lupus familiaris*) has evolved to display a remarkable phenotypic heterogeneity. From dachshund to grand danois, the current 400 dog breeds (Serpell & Duffy, 2014) manifest exceptional morphological, behavioral, and genetic diversity, providing valuable models for heredity, molecular genomics, and diseases (Wayne & Ostrander, 2007). Recent advances in long read sequencing have improved genome assemblies in various species, including human (Nurk *et al*., 2022) and dog (Jagannathan *et al*., 2021; Zhou *et al*., 2025). The refined annotations have enabled identification of ultraconserved elements (Snetkova *et al*., 2022), *loci* that influence evolution (Plassais *et al*., 2019; Vaysse *et al*., 2011; Sahlén *et al*., 2021), regions that produce capped RNAs (Hörtenhuber *et al*., 2024), and uncovered the dog’s epigenome in distinct tissues (Son *et al*., 2023). These analyses have compared DNA sequences between species and breeds, and measured stable RNAs. However, the functionality of a genome is largely manifested by the activity of its regulatory regions, which coordinate expression of protein-coding and non-coding (nc) RNAs. Transcription from a mammalian genome is considered pervasive (Clark *et al*., 2011), but only 1-3% of the DNA sequences encode for proteins (Palazzo and Lee, 2015). Indeed, 30% of human promoters direct synthesis of mRNAs (Nurk *et al*., 2022), and the various short and long ncRNAs carry versatile functions (Chadwik and Scott, 2013). Distal to promoters, enhancers (Banerji *et al*., 1981) coordinate differentiation (Herrmann et al., 2022), cell type-specific transcription (Wu and Huang, 2024), and stress responses (Himanen *et al*., 2022). Enhancers can recruit transcription factors and chromatin remodelers and produce unstable enhancer RNAs (eRNAs) in divergent orientations (Kim *et al*., 2010; Core *et al*., 2014). The characteristic pattern of engaged Pol II at genes and enhancers enables identification of functional genomic regions *de novo* (Danko *et al*., 2015; Rabenius *et al*., 2022), and predicts chromatin states with striking accuracy (Wang *et al*., 2022). Moreover, eRNA production correlates with the enhancer’s functional activity in cells and organisms (Tippens *et al*., 2018; Mikhaylichenko *et al*., 2018; Henriques *et al*., 2018; Carullo *et al*., 2020). Consequently, nascent RNA sequencing reveals active genes and enhancers in high sensitivity and resolution, quantifies enhancer activity, and tracks progression of Pol II complexes through the rate-limiting steps of transcription (Wissink *et al*., 2019; Vihervaara *et al*., 2023).

To investigate mechanisms of transcription at genes and enhancers, we analysed transcribing Pol II molecules in dog macrophage cells using Precision Run-On sequencing (PRO-seq; Kwak *et al*., 2013). PRO-seq adds a single biotinylated nucleotide to the 3’-ends of nascent RNAs, and subsequently, isolates the nascent transcripts with streptavidin. While the 3’-ends of PRO-seq reads report the active sites of transcription, the 5’-ends accumulate at the precise positions of Transcription Start Nucleotides (+1nts, TSNs; Vihervaara *et al*., 2023). In this study, identification of the +1nts and Pol II active sites uncovered architectures of initiation and pausing, pause distances, and sequences directing Pol II progression across dog genes and enhancers. To compare RNA synthesis to mature RNA expression, we performed strand-specific RNA-seq of polyadenylated (polyA+) transcripts. The matched data of synthesis and expression improved gene annotations, quantified RNA stability, and showed that genes predicted to encode ncRNAs produced unstable polyadenylated transcripts. To assay mechanisms that coordinate RNA synthesis and expression, we provoked the highly conserved heat shock response (HSR), which instantly reprograms gene and enhancer activity. Upon stress, transcription from coding and non-coding genes was coordinated *via* the rate-limiting step of Pol II pause-release. Instead, enhancer transcription was controlled through Pol II recruitment. For many genes, transcription remained unchanged and the mRNA expression upon stress was controlled *via* RNA stability.

Super-enhancers are clusters of transcribed enhancers and highly active distal regulatory elements that drive cell identity and cancer (Hnisz *et al*., 2013). Due to their association with key cellular processes and frequent mutations, super-enhancers are actively studied as potential drug targets and sources of genetic variation (Huang et al., 2018; Bacabac and Xu, 2023). Super-enhancers are traditionally identified as high Mediator 1 or H3K27ac enrichment in ChIP-seq, and the term indicating superiority without assessing functionality challenged (Pott and Lieb, 2015). Here, we identified clusters of transcribed enhancers (hereafter termed eClusters) in high precision and sensitivity from PRO-seq data, and analysed their enhancer content, transcriptional activity, and transcriptional regulation. The identified eClusters contained well-known super-enhancers, including the cancer-driving MYC and macrophage-activated EMB super-enhancers. We revealed the enhancer organisation and transcriptional activity of eClusters in detail, and found that clustered enhancers responded in unison, showing strikingly similar Pol II regulation upon acute stress. Enhancers within individual eClusters contained a higher sequence similarity than position or expression matched unclustered enhancers. However, their unified transcriptional response did not require the presence of the same DNA sequences. Taken together, this study provides a comprehensive characterization of transcription across genes and enhancers, including the precise architectures of initiation and pausing, Pol II progression through the rate-limiting steps of transcription, and mechanism of transcriptional reprogramming. We improve the calls of coding and ncRNAs in dog and measure RNA stability. Finally, eClusters and their enhancer activity is revealed, and their unified responses demonstrated. The precise coordinates and mechanisms of Pol II regulation provide a framework for functional annotations of various assays, including genome-wide association studies (GWAS), to discover regulatory principles driving RNA synthesis, genome-regulation, evolution, and complex traits.

## Results

### Nascent transcription and mRNA expression in dog macrophage cells

To quantify transcription across genes and enhancers, we performed PRO-seq in macrophage-monocyte DH82 cells, derived from golden retriever bone marrow (Wellman *et al*., 1988). The nascent transcription analyses were matched to strand-specific polyA+ RNA-seq (also called mRNA-seq) to measure both RNA synthesis and mature RNA expression (Fig. 1A-C). The sequenced reads were mapped to the most recent version of the dog genome, canFam6 (Jagannathan et al., 2021: Zhou *et al*., 2025). All the libraries showed high quality and reproducibility, as demonstrated with gene-by-gene correlations (rho > 0.95) and similar read distributions between biological replicates (Fig. S1). As exemplified by *β-actin* gene (*ATCB*), reads from polyA+ RNA-seq accumulated at exons, while the active sites of transcription deduced from PRO-seq were distributed across the gene (Fig. 1B). Unlike mature RNAs, transcription extended downstream of the gene into the termination window, and captured promoter-proximal Pol II pausing and divergent transcription, which are characteristic features of mammalian genes (Fig. 1B). Since PRO-seq detects synthesis of stable and unstable transcripts, it reports transcriptionally active genes and enhancers across the genome. We used discriminative regulatory element identification from global run-on data (dREG; Wang *et al*., 2019), to find the pattern of divergent transcription at promoters (Fig. 1B) and enhancers (Fig. 1C). We found 15,558 transcriptionally active genes among the 34,981 unique genes in the canFam6 genome, half of which produced detectable levels of polyA+ RNAs (Fig. 1D). In addition, a group of genes was expressed only at the polyA+ RNA level (Fig. 1D), indicating low RNA synthesis and stable mature RNAs. Distal to gene promoters, 42,396 putative enhancers were identified (Fig. 1D), defined as dREG-called sites of divergent transcription that occurred at least 1 kb away from any annotated transcription start site (TSS) of a gene (Fig. 1D). As visualized with heatmap, the putative enhancers showed the characteristic pattern of short divergent transcription (Fig. 1E).

**Figure 1.**
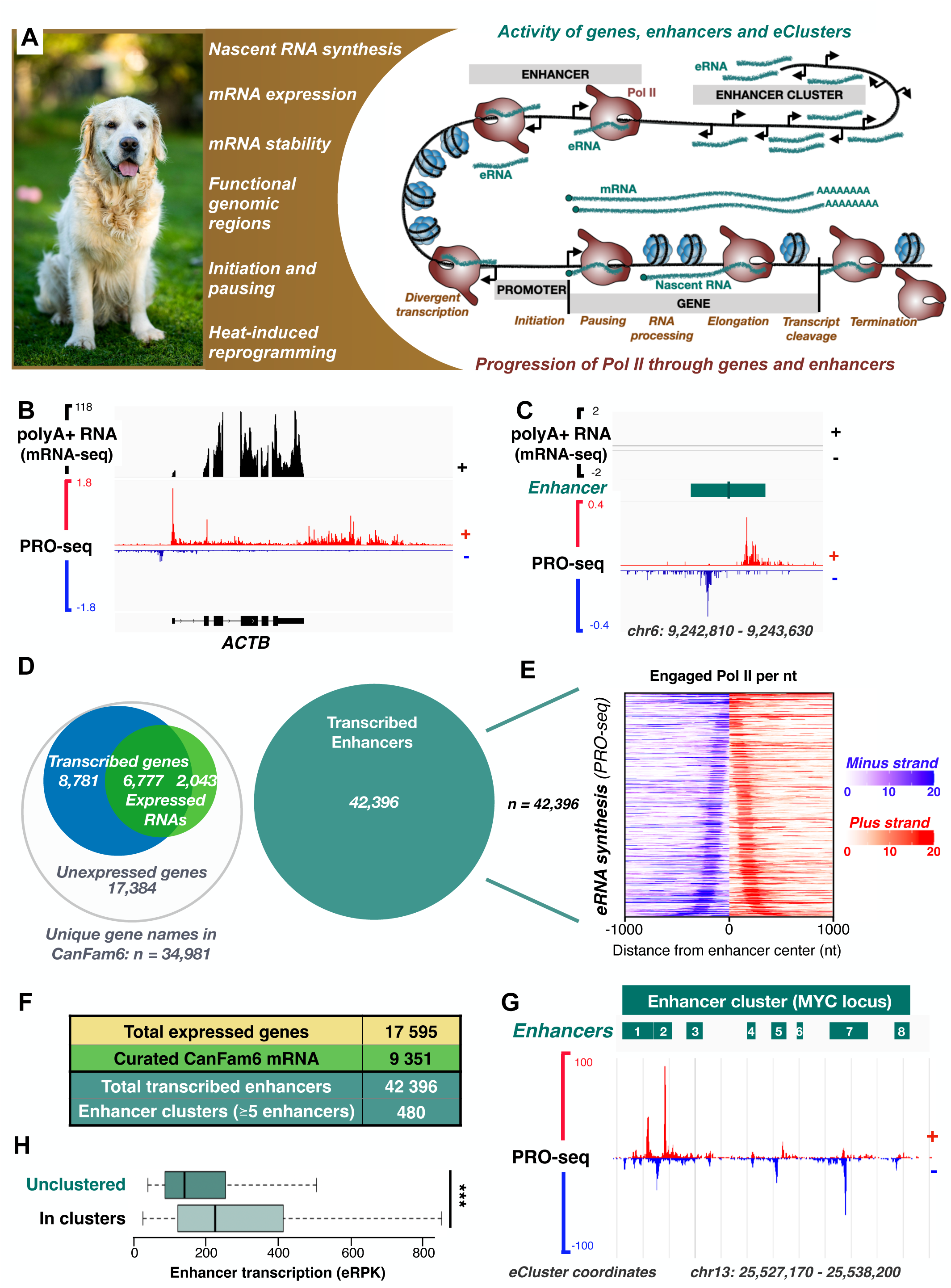
Quantifying RNA synthesis and mRNA expression across genes and enhancers in the dog genome. **A)** RNA synthesis and mRNA expression is analyzed in golden retriever macrophage (DH82) cells to i) identify transcribed genes and enhancers, ii) quantify RNA synthesis and mRNA expression, iii) measure mRNA stability, iv) uncover promoter and enhancer architecture, and v) characterize the transcriptional response to acute stress. **B)** Positions of engaged RNA Polymerases (PRO-seq) and mRNA expression (mRNA) of *b-actin* (*ACTB*) in DH82 cells. For mRNA (black), only the coding strand (+) is shown. For synthesis of nascent RNA, both coding (+ in red) and non-coding (- in blue) strands are visualized, revealing the characteristic pattern of divergent transcription from mammalian promoters. **C)** Genome browser example of a transcribed enhancer. The enhancer coordinates (teal), nascent transcription (PRO-seq) from + (red) and - (blue) strands, and the lack of mRNA (black) are shown. **D)** Left panel: Venn diagram comparing the number of genes producing nascent transcripts (azure blue), mRNA (leafy green), or neither (white). Right panel: Number of transcribed enhancers (teal) drawn to the scale of the genes shown in the Venn diagram. **E)** Heatmap quantifying enhancer RNA (eRNA) synthesis. eRNA produced from + strand is shown in red, and from - strand in blue. The shade of red or blue indicates engaged Pol II molecules per nucleotide (nt). **F)** Table summarizing the number of total expressed genes in DH82 cells (unison of transcribed and mRNA expressed), curated mRNA-coding genes (unison of mRNA-coding genes in CanFam6 refSeq + genes that show polyA+ RNA expression in DH82), transcribed enhancers in DH82 (divergent transcription distal to promoters), and number of enhancer clusters (>= 5 enhancers within 12,500 kb). **G)** Enhancer cluster on MYC locus, containing eight individual enhancers (e1-8). **H)** Quantification of eRNA synthesis at enhancers that do not locate into clusters (dark teal) or are part of enhancer clusters (light teal). The enhancer transcription is measured as density of engaged Pol II per kilobases of enhancer length (eRPK). The statistical difference was analyzed with ANOVA, ***p-value < 0.001.

Clusters of transcribed enhancers, also termed super-enhancers, contain several near-by enhancers that provide arrays of binding sites for transcription factors (Hniz et al., 2013; Whyte et al., 2013). We utilized the nucleotide-resolution profiles of engaged Pol II in PRO-seq data, and identified the precise locations of eClusters and their composition of individual enhancers (Fig. 1F-G). We found 480 eClusters in the dog macrophage cells, each including at least 5 individual enhancers within a 12.5 kb window (Fig. 1F). The initial 12.5 window was iteratively extended if additional enhancers resided within 2 kb, and the final eCluster location reported from the start of the first to the end of the last enhancer, capturing the precise coordinates of eClusters. One of the identified eClusters is a well-characterized super enhancer for *MYC* proto-oncogene, which in dog macrophage cells consisted of eight individual transcribed enhancers within a 11 kb region (Fig. 1G). Aligning with the initial studies (Hniz *et al*., 2013;

Whyte *et al*., 2013), the transcriptional activity of individual enhancers within an eCluster was higher than the activity of unclustered enhancers (Figs. 1H). This data is the first complete transcriptional profiling of the dog genome, involving nascent transcription of genes (Supplemental Dataset 1), enhancers (Supplemental Dataset 2), and eClusters (Supplemental Dataset 3), as well as expression levels of mature, polyA+ RNAs (Supplemental Dataset 4).

### Many genes predicted to encode ncRNA in CanFam6 produce unstable polyadenylated transcripts

CanFam6 genome includes annotations for genes that have been validated (n = 2,029) or predicted (n = 17,529) to encode mRNA (Fig. 2A-B). Of the validated mRNA genes, 1,051 were transcribed in dog macrophages, the vast majority of which (95%) also produced polyA+ RNA (Fig. 2B). Most transcribed genes in our data (n = 11,486) had been predicted to encode mRNA. Half of them (49%) expressed polyA+ RNA (Fig. 2B), albeit at a lower level than the validated mRNA genes (Fig. 2C). These results provide evidence for the expression of polyA+ mRNA from 6,115 genes, previously indicated to encode mRNA. The canFam6 also annotates genes that have been validated (n = 469) or predicted (n = 14,941) to encode ncRNAs (Fig. 2B). Intriguingly, large fractions of transcribed ncRNA genes expressed polyA+ RNA (Fig. 2A-B) at a low level (Fig. 2C). Polyadenylation is generally considered to select mRNAs. However, many ncRNAs are polyadenylated and spliced (Wilusz, 2016), which makes polyadenylation and exon visualization limited strategies for distinguishing mRNAs from ncRNAs. Our data demonstrates a clear correlation between high levels of polyA+ RNAs and validation for mRNA-coding. Indeed, projecting nascent transcription (PRO-seq) against mature RNA (polyA+ RNA- seq) demonstrated validated mRNA-coding genes to express high levels of polyA+ RNA from varying levels of transcription, indicating high RNA stability (Fig. 2D). In contrast, predicted mRNA genes encoded both stable and unstable RNAs, reflected in their wide ranges of transcription and RNA expression (Fig. 2D). In turn, predicted ncRNA genes produced unstable RNAs, as evidenced by their variation of transcription, and low levels of mature polyA+ RNA (Fig. 2D). These results imply that polyadenylated ncRNAs have a low half-life, or that gene predictions misclassify many unstable mRNAs as ncRNA.

**Figure 2.**
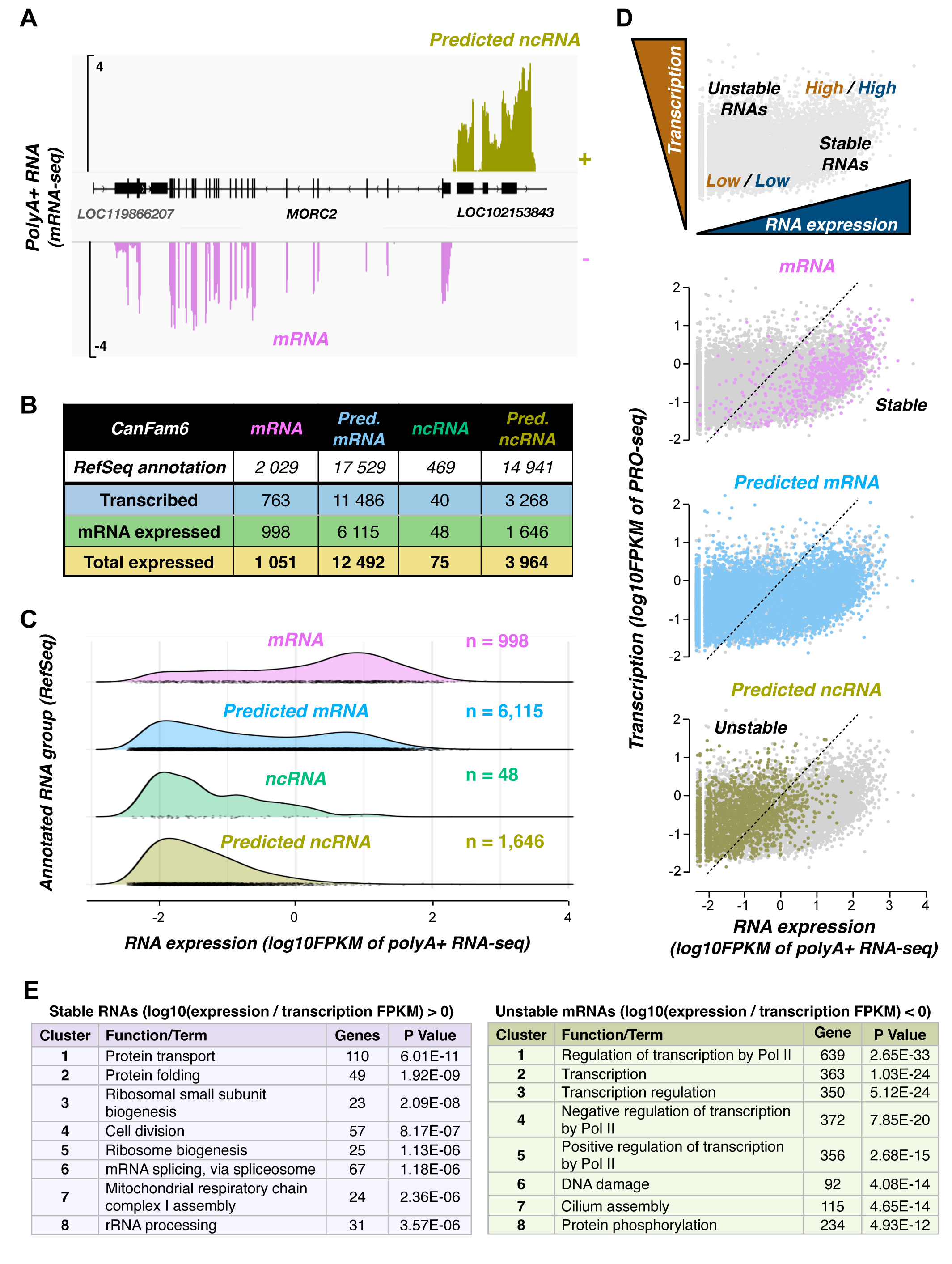
Identifying mRNA-coding genes and measuring mRNA stability. **A)** Strand-specific mRNA expression from genes that were categorized as mRNA-coding (*MORC2*) or predicted to be non-coding (*LOC102153843*) in CanFam6 RefSeq. **B)** Table counting transcribed and mRNA expressing genes, based on the category of the gene in RefSeq gene list (mRNA-coding, predicted mRNA-coding, ncRNA and predicted ncRNA). **C)** Quantification of mRNA expression according to the RefSeq category. **D)** Comparison of nascent transcription (engaged Pol II, y-axis) and mRNA expression (x-axis) from individual genes. High transcription with low mRNA expression indicates unstable RNAs, while low transcription with high mRNA expression indicate stable RNAs. Genes categorized as mRNA-coding by RefSeq produce stable mRNAs. Genes predicted to encode ncRNAs produce primarily unstable mRNAs. **E)** Gene ontology analysis displaying top eight biological functions associated with stable and unstable mRNAs.

We utilized the matched datasets of transcription and mature RNA expression to quantify RNA stability (polyA+ RNA-seq FPKM / PRO-seq FPKM). To identify cellular functions performed by the stable *versus* unstable transcripts, we ranked RNAs by their stability and used the DAVID annotation tool (Huang *et al*., 2009; Sherman *et al*., 2021) to call enriched molecular functions among the stable and unstable RNAs. Stable RNAs were associated with house-keeping functions, such as protein transport, protein folding and ribosome assembly (Fig. 2E). Instead, unstable RNAs were involved in control of transient processes, such as transcription regulation and protein phosphorylation (Fig. 2E). Taken together, matched RNA synthesis and expression quantified RNA stability and showed maintenance of core cellular processes to rely on highly stable RNAs, while unstable transcripts were associated with transient regulation, such as transcription control and signaling *via* post-translational modifications.

### Pol II progression across functional genomic regions in dog

Next, we identified functional genomic regions from the profile of active transcription, and investigated the progression of Pol II from the initiation, through pause-regulation, into productive elongation and termination (Figs. 3A-B). As exemplified by *ribosomal protein S9* gene (*RPS9*; Fig. 3A), PRO-seq quantifies engaged Pol II molecules at divergent transcription (purple), promoter-proximal pausing (orange), productive elongation (black), cleavage and polyadenylation site (CPS, light blue), and termination (pink). These measures provide means to investigate how Pol II proceeds through the stages of transcription. Genome-wide, the majority of active Pol II was involved in productive elongation across the gene bodies (Fig. 3B). Around 10 % of Pol II was engaged at pause-regions, termination windows, or enhancers, respectively (Fig. 3B). Only a small fraction of transcriptionally engaged Pol II was found at CPSs and sites of divergent transcription (Fig. 3B). Intriguingly, a third of total engaged Pol II localized to functionally unannotated regions. These regions included apparent unannotated genes (Fig. 3B) and genes where nascent transcription initiated tens of kilobases upstream the annotated TSS (Fig. S2A).

**Figure 3.**
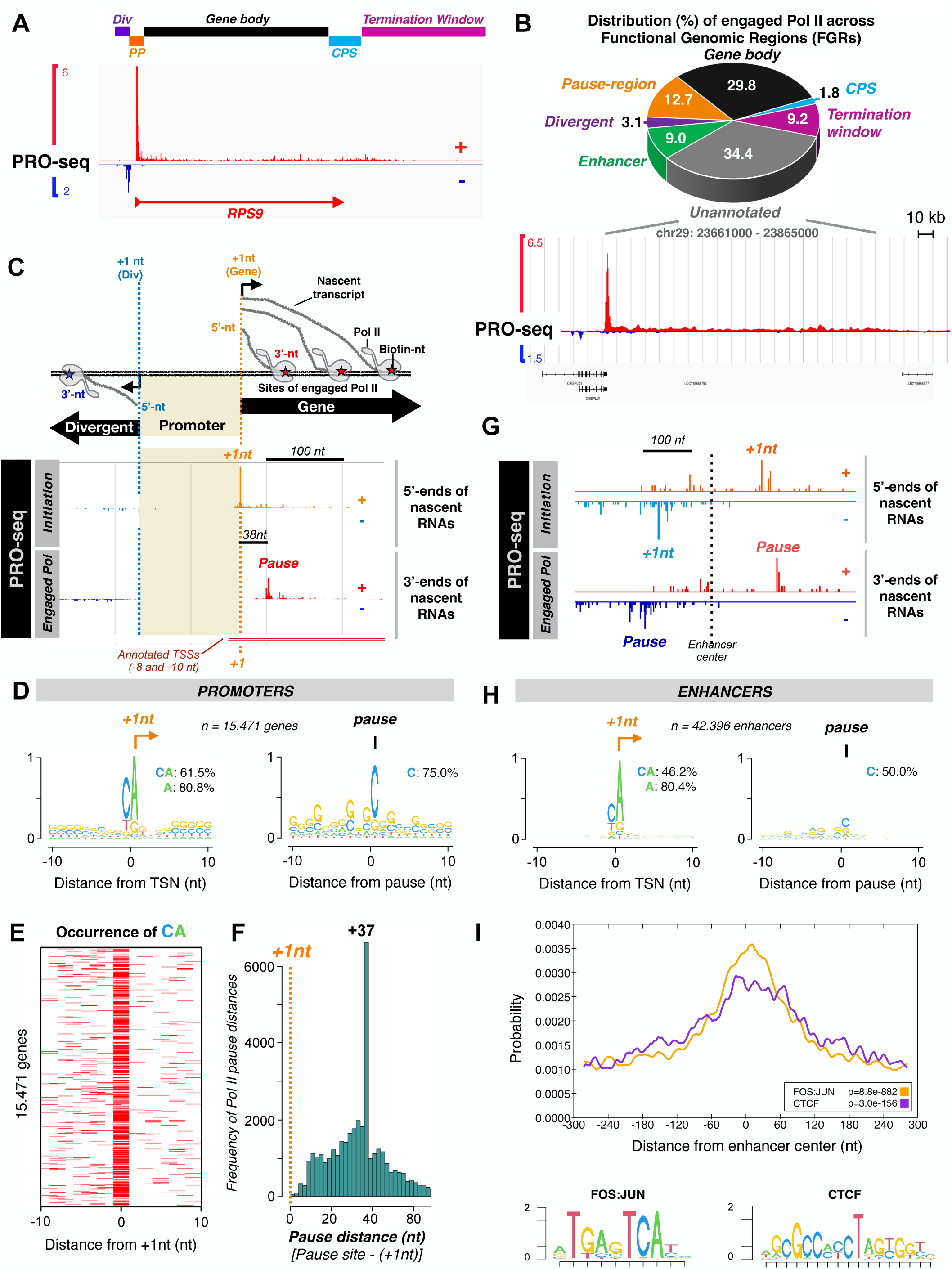
Functional genomic regions and transcriptional architecture of promoters and enhancers in the dog genome. **A)** Nascent transcription (PRO-seq) at *RPS9* gene. The profile of engaged Pol II reveals functional genomic regions, sites of divergent transcription (div), promoter-proximal region (PP), gene body, cleavage and polyadenylation site (CPS) and termination window. **B)** Distribution of engaged Pol II across the functional genomic regions. The browser track shows Pol II engagement across an unannotated genes, which shows sequence similarity to human ncRNAs. **C)** Transcriptional architecture of a promoter, showing the initiating nucleotides (+1 nt) toward the sense (gene) and anti-sense (divergent, div), the precise location of the promoter (shaded), and the pause coordinate (P) of Pol II on the gene. The +1 nts were identified from the 5’-ends of nascent transcripts, indicated on genome browser with S, and is compared to the annotated TSSs. **D)** Sequence enrichments at +-10 nt window around the +1nt (left) and Pol II pause coordinate (right), identified from all transcribed (n=15,558) genes. **E)** Heatmap of CA dinucleotide occurrence, showing the high enrichment of nucleotide C at the position -1nt, and nucleotide A at +1nt of transcribed genes. **F)** Histogram showing the distance of Pol II pausing from the precise +1 nt . **G)** Transcriptional architecture of a dog enhancer, showing the initiating nucleotides, pause nucleotides, enhancer center in-between. **H)** Sequence enrichments at +- 10 nt window around the +1nt (left) and Pol II pause coordinate (right), identified from transcribed enhancers. **I**) MEME-ChIP analysis at +-300 nt window around enhancer center, showing the distribution of FOS:JUN and CTCF motifs.

The single-nucleotide resolution of PRO-seq allows identification of the exact +1nt and the pause coordinate at each gene (Fig. 3C). Calling the +1nt to sense (gene) and anti-sense (divergent) directions revealed the precise positions of promoters, localizing between the divergently initiating Pol II complexes (Fig. 3C). Moreover, the +1nts and pause-coordinates allowed measurements of Pol II progression from the initiation through promoter-proximal pausing and into elongation (Figs. 3A and 3C). The coordinates of the annotated TSSs *versus* +1nts identified from the nascent transcripts can considerably differ (Figs. 3C and S2A). This discrepancy has been previously reported in fly (Nechaev *et al*., 2010), human (Tome *et al*., 2018; Vihervaara *et al*., 2023) and mosquito (van Hout *et al*., 2023). It is considered to arise from the traditional TSS calling relying on 5’- and 3’-end sequencing of total RNA, followed by reporting the longest gene coordinates to ensure capturing the whole transcript. Accordingly, the +1nt called from the nascent RNAs in dog localized downstream of the annotated TSS at majority of genes (Fig. S2B). Worth noting is that the annotation of TSSs from the total RNA was reliable for stable RNAs (Fig. S2C), however, a statistically significant decline in accuracy occurred with reduced RNA stability (Fig. S2D). We conclude that identification of the precise +1nts from nascent RNAs refines the promoter-calling, and is essential for detailed characterization of Pol II progression, transcription factor localization, and architectural analyses or promoters and enhancers.

### The architecture of initiation and pausing across dog promoters and enhancers

Using the precise +1nt and pause coordinates, we analysed the transcriptional architecture of promoters (Fig. 3C-F) and enhancers (Fig. 3G-I) in dog macrophage cells, including the locations of the initiator element (Inr) and sequences underlying Pol II pausing. Inr directs the initiation of transcription and is composed of the consensus sequence Py Py A_+1_ N T/A Py Py in mammals and fly (Javahery et al., 1994; Lo & Smale, 1996). We found Inr, heavily enriched for CA, at the +1nt of the vast majority of genes (Fig. 3D-E). Pol II pausing, instead, occurred at CG-rich regions, predominantly before incorporation of the cytosine (Fig. 3D), which is the least abundant ribonucleotide in mammalian cells (Traut, 1994). Next, we determined the pause-distance by subtracting the pause coordinate from the +1nt. Although we found varying pause-distances, a remarkable fraction of genes displayed the highest Pol II pausing at +37nt (Fig. 3F). This result is consistent with the previous studies showing Pol II pausing between 25- 60 nts downstream of the TSS in several model organisms (Rougvie and Lis, 1988, Nechaev *et al*., 2010; Kwak *et al*., 2013; Tome *et al*., 2018; Vihervaara *et al*., 2017). We predict that measuring pause-distances from the actual +1nt will narrow the range for Pol II pausing, and is essential for high-resolution analyses of Pol II dynamics at initiation, pausing, and pause-release across organisms.

Next, we investigated the architecture of enhancers. Since enhancers are divergently transcribed with comparable levels of Pol II initiation to both directions, two pairs of +1nts and pause coordinates were identified for each enhancer, one on the plus, the other on the minus strand (Fig. 3G). Analysis of the DNA sequences at enhancers’ +1nts revealed an identical Inr sequence that was found at the genes’ +1nt (Fig. 3H). Pol II undergoes pause-regulation also at enhancers (Henriques *et al*., 2018), although its engagement at enhancers is generally lower than at genes (Core *et al*., 2014). Like promoters, enhancers contained a GC-rich pause-region that decelerate Pol II, and a detectable enrichment of the C nucleotide downstream the pause (Fig. 3H). Furthermore, motif analysis showed that enhancer centers, localizing between the +1nts of divergently oriented Pol II complexes, were enriched for motifs of known enhancer binding proteins, including CCCTC-binding factor (CTCF) and activating protein 1 (AP-1), composed of Fos and Jun transcription factors (Fig. 3I). Altogether, we identified the exact coordinates of promoters (Fig. 3C) and enhancers (Fig. 3G) and deduced the genomic sequences underlying initiation and pausing. These coordinates are essential for correct positioning of mutations, transcription factor binding sites, regulatory elements, and chromatin connections, to analyse the molecular logic of genome-regulation in health, disease, and complex traits.

### Heat-induced reprogramming of dog genes occurs *via* Pol II pause-release

The HRS is an evolutionarily conserved mechanism that involves rapid induction of genes encoding heat shock proteins (HSPs) and their co-chaperones in response to proteotoxic stressors (Pessa *et al*., 2024). Since the HRS triggers an instant and profound change in the transcription (Vihervaara *et al*., 2018), it has become a commonly used model to investigate mechanisms of transcription. In this study, we characterized the HRS for the first time in dog by performing PRO-seq and polyA+ RNA-seq in DH82 cells exposed to 37°C and 42°C. For PRO-seq, a 30 min heat shock was used, whereas the polyA+ RNA-seq samples were collected at 30 min and 60 min of heat shock to monitor the slower changes in mature RNA levels.

Activation of HSR was observed across chaperone genes, such as the HSP family H member 1 (*HSPH1*), which showed a strong induction upon 30 min of heat stress (Fig. 4A). Besides activation, acute temperature stress triggers a prominent repression of transcription (Duarte *et al*., 2016; Mahat *et al*., 2016; Vihervaara *et al*., 2017), which in dog, was exemplified by the cyclin D1 gene (*CCND1*) (Fig. 4A). Altogether, heat shock triggered a transcriptional induction of 372 and a repression of 1,519 genes in dog macrophage cells (Fig. 4B). While heat-induced genes were related to stress responses and chaperone function, the repressed genes were highly enriched for cell cycle, cell division and protein phosphorylation (Fig. 4C).

**Figure 4.**
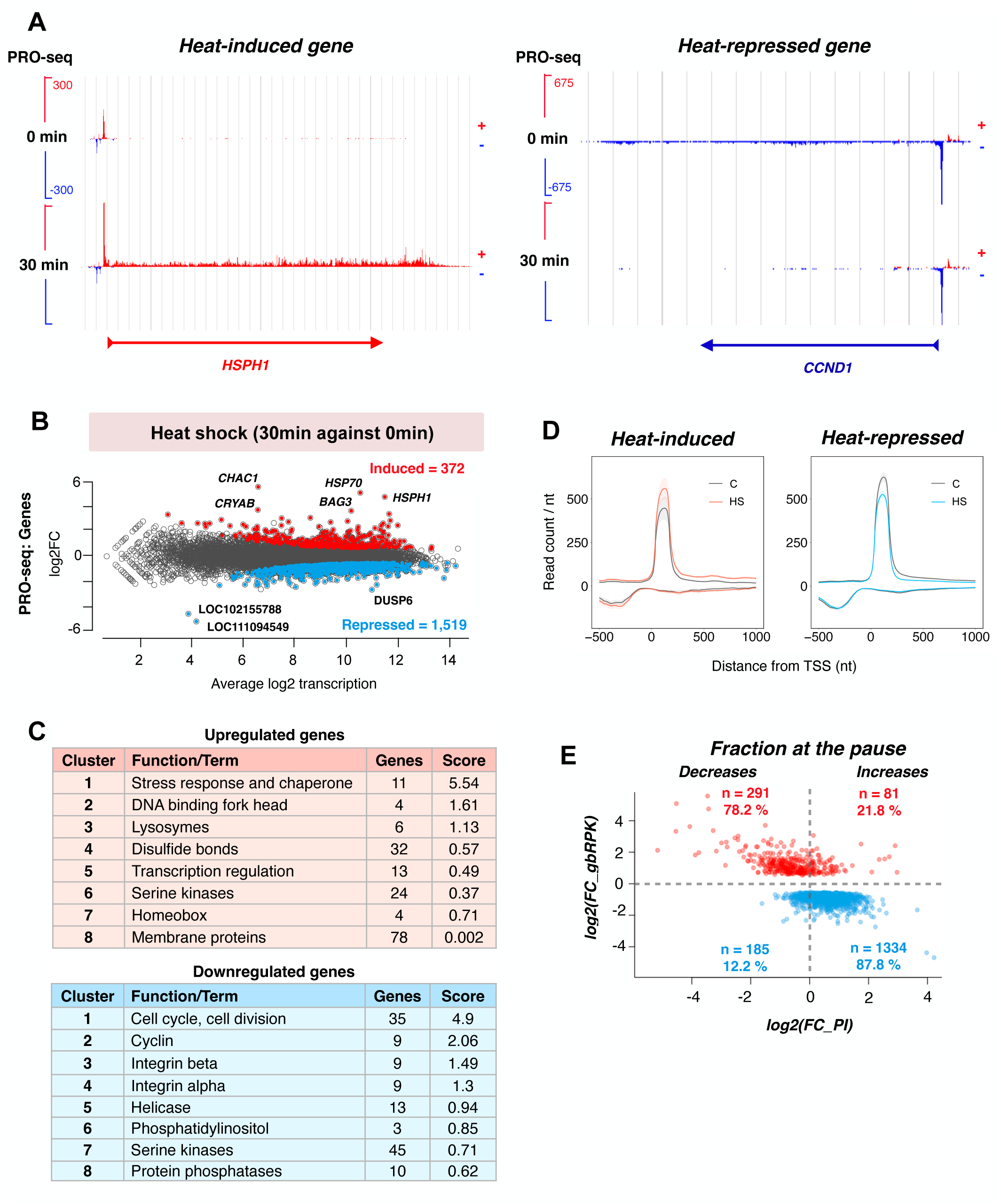
Promoter-proximal Pol II pause-release coordinates rapid transcriptional reprogramming. **A)** Heat-induced *HSPH1* (left) and heat-repressed *CCND1* (right) genes, showing the rapid change in nascent transcription upon acute stress. **B)** Genome-wide identification of significantly induced (n = 372; red) and repressed (n = 1,519; light blue) genes. The MA-plot is generated with spike-in scaled and DESeq2 analyzed differential expression using p-value < 0.05 and absolute fold change > 1.25. **C)** Gene Ontology analyses of enriched function among the induced (red table) and repressed (light blue table). The top eight clusters are shown. **D)** Average density of engaged Pol II at promoter-proximal regions of heat-induced (left) and heat-repressed (right) genes. **E)** The change in Pol II pausing index (PI), as a function of heat-induced change in the RNA synthesis at each induced or repressed gene. The percentage of induced (red) and repressed (light blue) genes that decrease (left) versus increase (right) Pol II pausing upon heat shock are indicated.

Earlier studies have identified promoter-proximal pause-release as the key step that regulates both the induction and repression of transcription (Vihervaara *et al*., 2018; Core and Adelman, 2019). To investigate mechanisms that reprogram transcription in dog cells, we measured engaged Pol II molecules within promoter-proximal regions and across gene bodies. Heat-induced genes gained both paused and elongating Pol II, which demonstrates accelerated recruitment, coupled to efficient release of Pol II into productive elongation (Figs. 4A and 4D). Instead, the repressed genes showed an opposite pattern, maintaining Pol II at the pause and preventing its entry into elongation (Figs. 4B and 4D). Consequently, the pausing index, which is the ratio of paused Pol II per elongating Pol II, decreased at heat-induced genes (Fig. 4E), evidencing an increased rate of Pol II entry into productive elongation. Instead, the pausing index at repressed genes elevated upon heat shock due to blocking Pol II at the pause (Fig. 4E). We conclude that HSR across dog genes is coordinated at the promoter-proximal pause-regulation, activating pause-release at chaperone genes, and inhibiting pause-release at cell cycle genes.

### Heat shock changes transcription and RNA stability

Upon heat shock, differentially expressed mature RNAs were detected upon a 30-min heat shock, reaching a total of 233 upregulated and 38 downregulated polyA+ RNAs upon 60 minutes (Fig. 5A). For many mature RNAs, the initial change at 30 min was amplified at 60 min, as demonstrated with *HSPH1* and *CCND1* (Fig. 5B). To understand kinetics of RNA synthesis and mature RNA expression, we compared PRO-seq and polyA+ RNA-seq signals during heat shock. As expected, RNA synthesis showed faster and more prominent changes than mature RNA expression, which is particularly evident for the repressed genes (Fig. 5C-D). Indeed, detecting reduction in mature RNA requires degradation of the existing RNAs. Likewise, detecting increased levels requires production of sufficient quantities of new over the existing RNAs. Together, the matched PRO-seq and polyA+ RNA-seq allowed analysing changes that occurred at the levels of transcription and expression (Fig. 5C). These analyses revealed RNAs that were induced or repressed *via* transcription, including *BAG3* and *DUSP6*, or *via* stability, including *ATF4* and *CCL8* (Fig. 5C). Altogether, our data reports rapid heat-induced changes in RNA synthesis and RNA expression, and distinguishes whether the expression changes are provoked *via* transcriptional regulation or changed RNA stability.

**Figure 5.**
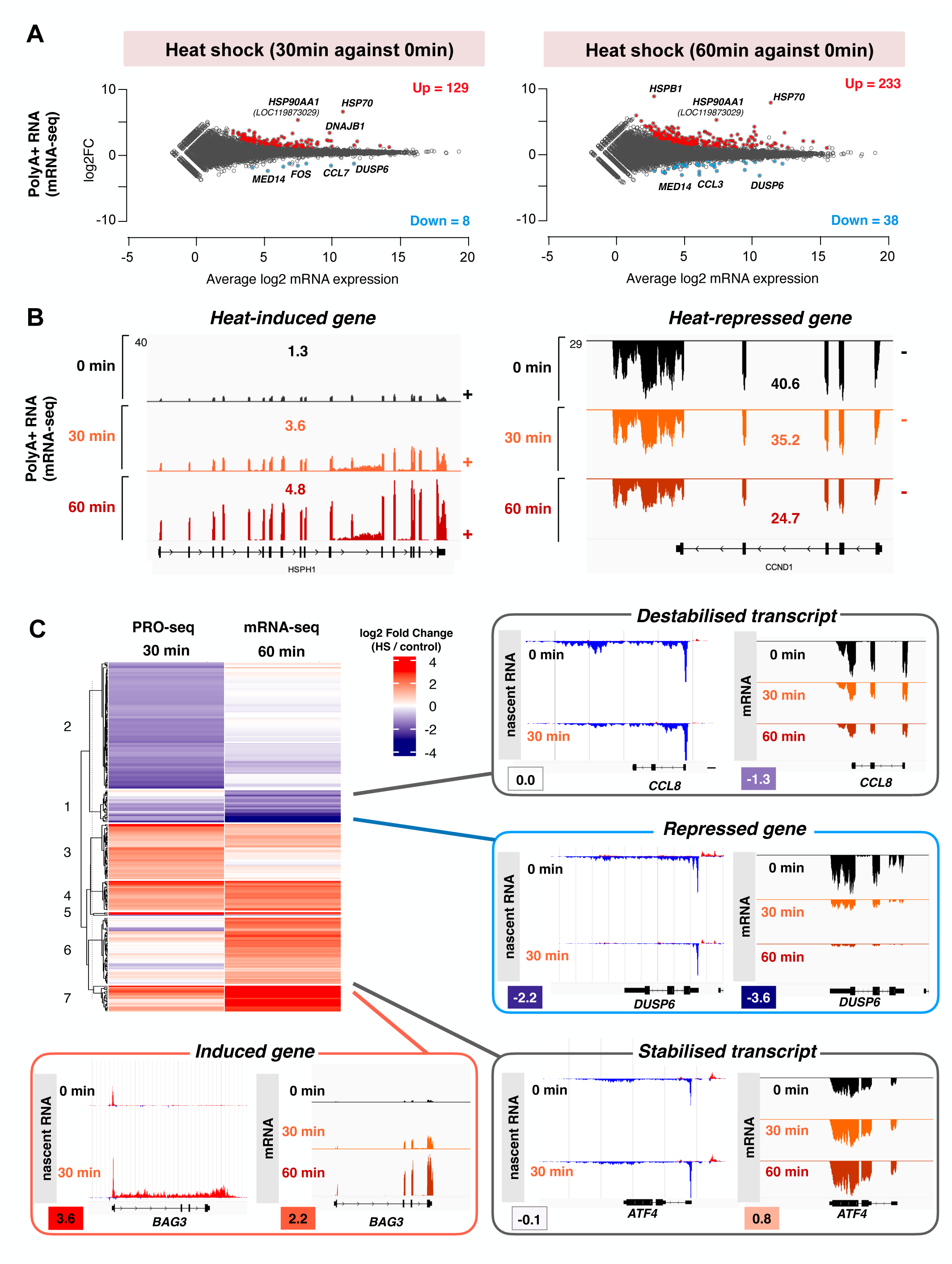
The rapid reprogramming of nascent transcription is followed by slowly changing mRNA expression. **A)** Genome-wide identification of mRNAs with significantly increased (red) or decreased (light blue) expression upon heat shock. MA-plots of spike-in scaled and DESeq2 analyzed differential expression of mRNAs using p-value < 0.05 and absolute fold change > 1.25. **B)** Genome browser profiles of mRNA expression from the transcriptionally activated *HSPH1* (left) and repressed *CCND1* (right) genes. **C)** Heat-induced change of nascent transcription (PRO-seq) and mRNA expression (mRNA-seq) compared across the dog genes with heatmap. Four example genes are illustrated in genome browser, indicating transcriptional induction (*BAG3*), transcriptional repression (*DUSP6*), as well as mRNAs whose levels change without transcriptional changes *via* increased (*ATF4*) or decreased (*CCL8*) stability.

### Distinct mechanisms reprogram transcription at promoters and enhancers

One of the benefits of tracking engaged Pol II molecules is the ability to measure changes in the synthesis of unstable RNAs, including eRNAs. Exposure to 30 min heat shock increased Pol II engagement at 796, and reduced Pol II engagement, at 229 enhancers (Fig. 6A). Unlike genes, which were primarily induced and repressed through Pol II pause-regulation, the enhancer transcription was coordinated *via* initiation (Fig. 6B-C). Particularly, the induced enhancers recruited Pol II in response to heat shock, gaining a higher Pol II density at and downstream of the enhancer pause (Fig. 6B-C). Downregulated enhancers, in turn, showed Pol II pausing under normal growth conditions, and lost Pol II from the whole enhancer when exposed to heat shock (Figs. 6B and 6D). These results identify striking differences in the molecular mechanisms that reprogram Pol II at genes *versus* enhancers. Despite the distinct mechanisms, genes and enhancers that resided in close proximity showed a similar direction of response, either gaining or losing Pol II. The concordant induction of a gene and its nearby enhancer is exemplified by *serine hydroxymethyltransferase 2* (*SHTM2*) *locus* (Fig. 6C), and the concordant repression of a gene-enhancer pair with *MTSS I-BAR domain containing 1* (*MTSS1*) and its upstream enhancer (Fig. 6D).

**Figure 6.**
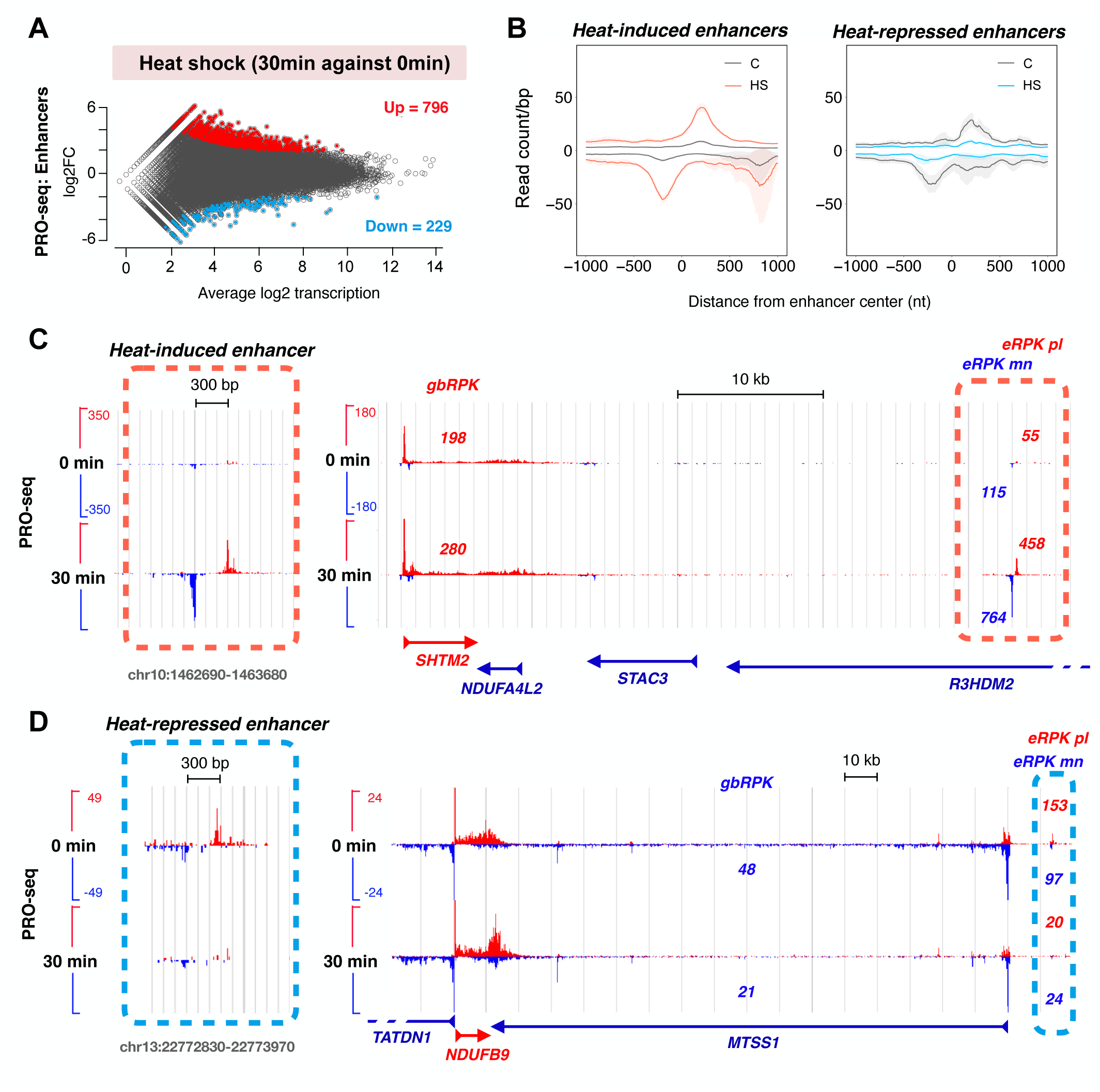
Heat-induced reprogramming of enhancer transcription. **A)** Genome-wide identification of enhancers with significantly induced (n = 796; red) or repressed (n = 229; light blue) eRNA synthesis upon heat shock. MA-plot of spike-in scaled and DESeq2 analyzed differential expression using p-value < 0.05 and absolute fold change > 1.25. **B)** Average density of engaged Pol II at heat-induced (left) and heat-repressed (right) enhancers. **C)** Example of an heat-induced enhancer, and its localization close to *SHTM2* gene that gains Pol II upon heat shock. **D)** Example of an heat-repressed enhancer, and its localization close to *MTSS1* gene that loses Pol II upon heat shock.

### Enhancers within an eCluster mount a unified response

The concordant response of near-by genes and enhancers prompted us to analyse transcriptional responses within eClusters. First, we addressed whether clustering gives an enhancer a larger gain or loss of Pol II upon transcriptional reprogramming (Fig. 7A). Intriguingly, the most prominent gains and losses of Pol II were observed at enhancers that displayed high basal transcription, regardless of whether the enhancer was located in a cluster or not (Fig. 7A). This data shows that clustering itself does not change the magnitude of an enhancer’s heat-responsiveness. However, the average basal transcription of a clustered enhancer is higher than that of an unclustered enhancer (Fig. 1H), which positioned the clustered enhancers toward the end of steeper changes (Fig. 7A). Next, we analyzed the heat-responsiveness of enhancers that were clustered together. We found that the vast majority of enhancers within an eCluster were regulated to the same direction, undergoing either transcriptional induction (Fig. 7B) or reduction (Fig. 7C). Analyses of all eClusters revealed a strong unified response, as enhancers within an eCluster showed the same direction of response (Fig. 7D) with binomial test p-value < 2.2*10^-16^ against expected similarity from all enhancers (Fig. S3A). To quantify whether the unified response was contained within the eClusters, we generated two matched controls from unclustered enhancers (Fig. S3B). First, we selected position-matched unclustered enhancers from the vicinity of each eCluster, generating the same number of control groups and individual enhancers therein. Second, we used the R sample package and formed groups of unclustered enhancers with a matching enhancer count and eRNA production to eClusters (Fig. S3B). Compared to the control groups, eClusters showed substantially greater similarity in enhancer responses, indicated by the fraction of enhancers within a group showing the same direction of change (Fig. 7E), and statistical comparisons between the eClusters and the matched controls (Fig. S3A,C-D).

**Figure 7.**
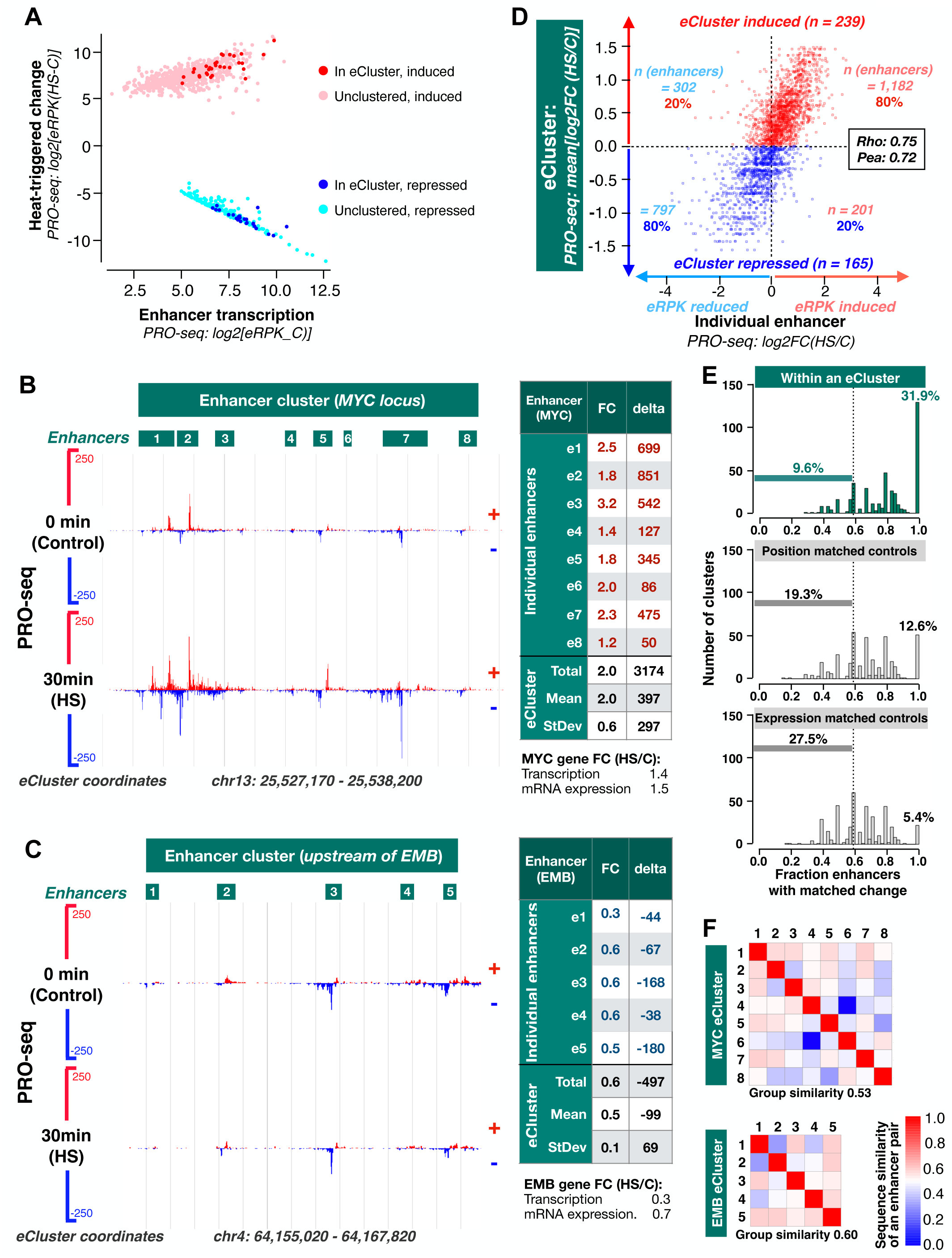
Clustered enhancers respond in unison. **A)** Comparison of reads per kilobase log2(eRPK) and change in eRPK upon heat shock log2(eRPK, HS-C) among clustered and unclustered enhancers. Clustered enhancers (eClusters, also known as super-enhancers) contain a minimum of five enhancers within a 12.5 kb window. **B-C)** Left: Transcriptional response of individual enhancers within the MYC eCluster (B) and EMB eCluster (C). The coordinates of the eCluster and its individual enhancers are shown. Right: Table listing the heat-induced fold change (FC, HS/C) and absolute change (delta, HC-C) of eRPK within each enhancer. Increased eRPK is shown in red, reduced eRPK in blue. The transcriptional change (PRO-seq and mRNA-seq) of the MYC (B) and EMB (C) genes are shown below the respective table. **D)** The transcriptional change (log2FC, HS/C) of individual enhancers within an eCluster, compared to eCluster’s total change [mean(log2FC, HS/C). Induced eClusters are show in red, repressed eCluster in blue. Individual enhancers within an eCluster for a horizontal row. The top right quadrant contains induced enhancers within induced eClusters. The bottom left quadrant contains repressed enhancers within repressed eClusters. The top left and bottom right contain enhancers that do not follow the direction of the eCluster’s transcriptional change. **E)** Comparison of fraction clustered enhancers that show the same directionality of change as the eCluster. The enhancers within eClusters (top, teal) are compared to position matched (middle, gray) and expression matched (bottom, grey) unclustered enhancers. **F)** Correlation matrix showing pairwise comparisons of sequence similarities between enhancers of the MYC eCluster (top) and the EMB (bottom) eCluster. The color scale (red-white-blue) indicates the sequence similarities, ranging from all k-mers shared (red) no k-mers shared (blue). Group similarity score measures similarity of all pairwise comparisons within the respective eCluster.

Enhancer clustering is considered to increase the binding opportunities for transcription factors, enabling the cluster’s responsiveness to several stimuli and providing regulatory variability. We asked whether the similar regulation of Pol II was coordinated by co-occurring DNA sequences within enhancers of the same eCluster. First, we used mash sketch (Ondov *et al*., 2016) to identify all possible 9-mer sequences at each enhancer, and next, quantified pairwise the k-mer similarity between enhancers. As compared to the position matched and expression matched controls, enhancers within an eCluster showed a subtle but statistically significant increase in k-mer similarity (Fig. S3E). This sequence similarity could contribute to the process of clustering, that is, to provide functional logic for the formation of an eCluster. Alternatively, the sequence similarity could be the basis for the similar enhancer responses. Detailed analyses of k-mers within all the eClusters revealed that clustered enhancers did not need sequence similarity for a unified response (Figs. 7F and S4). As an example, enhancers 2 and 8 within the MYC eCluster (Fig. 7B and 7F), and enhancers 2 and 3 within the EMB eCluster (Fig. 7C and 7F), shared only two and one 9-mers, respectively, out of over 1000 distinct 9- mers identified. In accordance, highly varying k-mer similarities (Fig. S4A) we detected at the distinct eClusters, including compartmentalization of similarity (Fig. S4B). Together, these results reveal a pervasive heat-induced reprogramming of enhancer transcription in dog cells, with coordinated changes of enhancer activity within eClusters. In summary, our study provides the first comprehensive characterization of dog transcriptome, heat shock response, and the architecture of promoters, enhancers, and eClusters, and the unified responses of connected regulatory elements.

## Discussion

### Characterization of dog transcriptome at single-nucleotide resolution

Identification of transcriptionally active regulatory elements and their precise locations is crucial for functional interpretation of genome analyses, including GWAS and molecular genomics. *Canis lupus familiaris* is phenotypically highly diverse, consisting of 400 distinct breeds (Serpell & Duffy, 2014), and together with other canine species, makes dog an excellent model for analysis of disease linkages and molecular mechanisms of evolution, complex traits, and genome-regulation. To facilitate genome research, several projects have assayed regulatory regions in dogs *via* DNA sequences and stable RNA expression (Hoeppner *et al*., 2014; Wang *et al*., 2021). Additionally, consortia such as the dog genome annotation project (DoGA), have important roles in filling in missing gaps in the annotations of genes and other genomic regions (https://www.doggenomeannotation.org/). Using PRO-seq, we tracked engaged Pol II molecules at nucleotide-resolution across the dog genome (Figs.1 and 3). These analyses tackled for the first time how the process of transcription across genes and enhancers is coordinated in dog cells. With the matched polyA+ RNA-seq data, the profiles of engaged Pol II provide tools for improved annotations of coding and non-coding genes (Figs. 3B and S2A). Moreover, the positions and transcriptional architectures of genes, enhancers and eClusters (Figs. 2, 6-7), provide functional annotations yet missing in the dog. Currently, the dog genome annotations remain incomplete, which is reflected in the 30% fraction of engaged Pol II molecules at unannotated regions (Fig. 2B), as compared to 9% in the human genome (Rabenius *et al*., 2022). To facilitate detailed functional annotation, we have assembled the coordinates of promoters, enhancers and eClusters, as well as polyA+ RNA expression and RNA stability in Supplementary Datasets S1-4. The raw data and normalized coverage file have been deposited to Gene Expression Omnibus. Our analyses also provide the positions of transcription initiation and pause coordinates, which we propose can be used as a framework for GWAS analyses to assay precise positions of mutations or single nucleotide polymorphisms (SNPs) with respect to the exact coordinates of Pol II regulation, including distances to transcription initiation, Pol II pausing, productive elongation, and termination.

### Bridging GWAS to mechanisms of transcription

Single-nucleotide resolution of PRO-seq allows assigning mutations and SNPs in the genome to regulation of RNA polymerases. In this study, we used heat shock as a model to analyze Pol II control across the genome, including recruitment, pause release, elongation, divergent transcription, and changes in enhancer activity. Similarly, PRO-seq can be utilized to uncover how distinct processes of transcription at genes and other regulatory regions are impacted by diseases, disease-associated mutations, and mutations that play a critical role in evolution. This important implication allows linking mutations to not only gene expression changes but also to mechanisms of Pol II regulation. In the future, directed mutations combined to functional analyses with nascent RNA sequencing will aid in deducing the molecular logic behind complex traits. Furthermore, by combining PRO-seq with polyA+ or total RNA-seq, transcription mechanisms can be linked to RNA expression and stability, allowing comprehensive gene expression profiling.

### Detailed architecture of enhancer clusters

Previous studies have found that clustered enhancers increase the binding site repertoire for regulatory proteins. Moreover, the clustered enhancers can regulate target genes additively, although the impact of each enhancer varies (Hnisz et al., 2015; Hay et al., 2016). Recently, the locus control region for alpha globin genes was used as a model for a super-enhancer, and shown to contain regular enhancers as well as facilitators (Blayney et al., 2023). To assay the functional impact of its elements, distinct enhancers within super-enhancers need to be identified. Current means to identify, and name, super-enhancers have provoked discussion as enrichment of Mediator or H3K27ac does not provide a direct measure of functionality, or detailed architecture of its enhancers. Moreover, ChIP-seq signals are broad and the thresholds for enhancer count and region length somewhat arbitrary. To date, the term super-enhancer has been widely used for various types of regulatory regions, including locus control regions for alpha- and beta-globin genes (Grosveld et al., 1987; Jarman et al., 1991), leaving the concept of super-enhancers ambiguous. In this study, we leveraged the ability of PRO-seq to identify transcribed enhancers in high resolution and sensitivity, and developed a program, eClusterer, that finds clusters of transcribed enhancers and reports their precise architecture and transcriptional activity. The eClusterer provides one read-out for functionality, the activity of eRNA production, but like the traditional super-enhancer calling, it does not give a direct measure of functional activity regarding gene regulation. Taken together, eClusterer enables high-resolution analysis of clustered enhancers, and measures of eRNA synthesis from its individual members, which can be highly informative for discovering the architectural principles and regulatory logic of enhancers that work in unison.

### Coordinated transcription across genes and enhancers

HSR provides an excellent model to assay mechanism of transcriptional reprogramming. Using heat shock as a model, we show that dog cells rapidly induced hundred and repressed over a thousand genes *via* Pol II pause-regulation (Fig. 4). Instead, the transcriptional response at enhancers was coordinated *via* Pol II recruitment and initiation (Figure 6). Despite the distinct step of Pol II regulation, linked genes and enhancers responded in similar manner, either increasing or decreasing transcriptional activity (Figure 6). Moreover, eClusters were shown to mount a unified response, displaying either a simultaneous induction or repression of individual enhancers (Fig. 7). Although clustered enhancers displayed a higher sequence similarity than unclustered enhancers (Fig. 7), their similar transcriptional response did not need the presence or the same 9-mer sequences. Overall, the increased sequence similarity could provide a framework for the enhancer clustering, and the strong transcriptional signals occurring within one of the enhancers affect the whole eCluster, possibly also its connected genes. Taken together, the data presented in this study is the first complete nucleotide-resolution characterization of the enhancer landscape in dog, involving +1nt and pause coordinates, their sequence composition, and the coordinates of eClusters. Future studies hold exciting potential for dissecting the molecular mechanisms driving the unified response of clustered enhancers and the impact of transcription signal over the enhancers and facilitators within an eCluster.

### Conclusions

Here, we used a combination of PRO-seq and mRNA-seq to perform a comprehensive single-nucleotide-resolution characterization of dog transcriptome, including measurement of engaged Pol II in pause regions, gene bodies, termination sites, enhancers, enhancer clusters and quantification of RNA stability. Moreover, we provide coordinates for enhancer clusters and +1nt and pause nucleotide coordinates for both transcribed genes and enhancers, which can be used as a framework for future GWAS studies. In future studies, it will be important to investigate how distinct mutations impact mechanisms of transcription, including initiation, pause release, and progression of RNA polymerases across genes and enhancers. Bridging genetic variants associated with behavioural traits and diseases to mechanisms of transcription will have an important impact on human medicine, allowing a better understanding of transcriptional changes observed in various diseases.

## Material and Methods

### Cell culture and heat shock treatment

The dog macrophage-monocyte DH82 cell line (Wellman *et al*., 1988) was obtained from ATCC (CRL-3590) and cultured in EMEM supplemented with 10 % fetal bovine serum, 0.292 g/l l-glutamine, 50 µg/ml streptomycin/penicillin, and non-essential amino acids (Gibco). The cells were maintained in a humidified 5% CO_2_ atmosphere at 37°C. Heat shock treatments were performed by submerging cells into a water bath prewarmed to 42°C. For PRO-seq experiments, 20 million cells seeded to 150 mm plates in 15 ml media were exposed to a heat shock for 30 min, whereas for RNA-seq, 2 million cells seeded to 6-well plates in 2 ml media were exposed to a heat shock for 30 or 60 min.

### PRO-seq

PRO-seq was performed as previously described (Mahat *et al*., 2016; Vihervaara *et al*., 2023) from two biological replicates using DH82 cells which were either untreated (37°C) or exposed to a 30 min heat shock at 42°C. Cells were washed with ice-cold PBS and chromatin isolated by resuspending the pelleted cells in ice-cold NUN buffer (0.3 M NaCl, 1 M Urea, 1% NP-40, 20 mM HEPES pH 7.5, 7.5 mM MgCl, 0.2 mM EDTA, 1 mM DTT, 20 U SUPERaseIn RNA inhibitor [Thermo Fisher], 1x cOmplete EDTA-free protease inhibitor cocktail [Roche]). Samples were vortexed and centrifuged (12,500 g, 30 min, 4°C). The chromatin pellet was washed with 50 mM Tris-HCl (pH 7.5) and resuspended in chromatin storage buffer (50 mM Tris-HCl pH 8.0, 25% glycerol, 5 mM MgAc_2_, 0.1 mM EDTA, 5 mM DTT, 20 U SUPERaseIn RNase Inhibitor). The chromatin was sonicated for 10 min with 30 s on/off intervals using sonicator by Bioruptor (Diagenode).

Before run-on reactions, spike-in chromatin from *Drosophila* S2 cells was added to samples, counted to constitute 1% of the total chromatin. Run-on reactions were performed for 5 min at 37°C using biotinylated nucleotides (5 mM Tris-HCl pH 8.0, 2.5 mM MgCl_2_, 150 mM KCl, 0.5 mM DTT, 0.5% Sarkosyl, 20 mM biotin-ATP/GTP/CTP/UTP [Perkin Elmer], 20 U SUPERaseIn RNA inhibitor). Total RNA was isolated with Trizol followed by ethanol precipitation. Base hydrolysis with NaOH was used to fragment the RNA, after which samples were purified from unincorporated biotin-NTPs using P-30 columns (Bio-Rad). Nascent RNAs were isolated with streptavidin C1 beads (Dynabeads MyOne Streptavidin C1, Thermo Fisher) and purified using Trizol and ethanol precipitation. Libraries were generated with Tru-Seq small-RNA adaptors, starting with the ligation of the 3’ adaptor, followed by an additional isolation with streptavidin C1 beads. 5’-cap was removed from nascent RNAs using RNA 5’ pyrophosphohydrolase (RppH, NEB), after which 5’ ends were phosphorylated with T4 polynucleotide kinase (NEB) and 5’ adapter was ligated. Nascent RNA was isolated with streptavidin C1 beads, followed by reverse transcription, PCR amplification, purification of the final libraries with Mag-Bind TotalPure NGS magnetic beads (Omega Bio-Tek), and sequencing using NovaSeq 6000. The sequencing was conducted by National Genomics Infrastructure (NGI), Sweden. The raw reads and normalized coverage files have been deposited to Gene Expression Omnibus (GEO).

### PolyA+ RNA-seq

Libraries for polyA+ RNA-seq were prepared with Illumina Stranded mRNA Prep kit (Illumina) from 1 mg total RNA as starting material. The PolyA+ RNA-seq was performed from three biological replicates of DH82 cells, exposed to either 0 min (control), 30 min or 60 min heat shock at 42°C. Total RNA was isolated from cells using Trizol, followed by ethanol precipitation. PolyA+ RNA was selected as instructed in Illuminas Stranded mRNA Prep Kit, and converted to cDNA using reverse transcription with random primers. Adapters were ligated to the ends of cDNA, and the libraries were amplified with PCR, purified and sequenced by national Genomics Infrastructure (NGI) Sweden, using NovaSeq 6000. The raw reads and normalized coverage files have been deposited to GEO.

### Computational analyses

#### Analysis of mRNA-seq data

Sequencing reads were mapped to the dog genome (canFam6) using HISAT2 (Kim *et al*., 2019) with parameters: -k 10 --rna-strandness FR --no-mixed --no-discordant. The level of mRNA expression was quantified from exons, using FeatureCounts (Liao *et al*., 2014) with parameters: -p -B -s 2. The output files from FeatureCounts were used in DESeq2 (Love *et al*., 2014) to perform differential gene expression analysis. To call statistically significant changes, adjusted p-value threshold of 0.001 and absolute fold change threshold of 1.25 were used. Sequencing depth normalization was used for the normalization of the mRNA-seq data.

#### Analysis of PRO-seq data

PRO-seq adapters from paired-end data were removed using the FastP tool (Chen 2023). The sequence of the read 1 was reverse complemented using Fastx toolkit (https://github.com/agordon/fastx_toolkit). Sequenced reads were mapped to the dog genome (canFam6) using Bowtie2 (Langmead & Salzburg, 2012) with parameters: -q --end-to-end –ff. SAM file output from Bowtie2 was converted to BED format. From the BED files, the most 5’ and 3’ single-nucleotide coordinates were retained separately for reads originating from plus and minus strands. These single-nucleotide coordinates were used to identify transcription the precise +1nts, and pause coordinates at single-nucleotide resolution as described in Vihervaara et al., 2023.

### Normalization of PRO-seq data

PRO-seq data was normalized using read counts in the ends of long genes, which has provided a robust normalization strategy in previous studies (Vihervaara *et al*., 2017). Normalization region was defined as +100 kb from TSS to -0.5 kb from the CPS of genes that were over 150 kb in length. Since Pol II moves 3 kb/min at most genes (Fuchs et al., 2014; Jonkers et al., 2014), advancing and receding waves of Pol II triggered by 30 min heat shock will not have enough time to reach the normalization region.

### Quantification of gene transcription

Actively transcribed genes were identified using dREG (Wang et al., 2019), which detects sites of divergent transcription from PRO-seq data, identifying both genes and enhancers. dREG coordinates were determined separately in each sample, after which the dREG coordinates were merged into a single file containing transcription initiation sites across all samples. dREG coordinates were then intersected with the annotated TSSs in canFam6. Pol II pausing was quantified from a 50 nt window, centered on the highest coordinate of Pol II engagement (pause nucleotide). Transcriptional activity of a gene was quantified by counting reads per kilobase DNA within gene bodies (+500 from TSS to -500 from CPS).

### Quantification of enhancer transcription

Actively transcribed enhancers were identified as dREG-detects sites of divergent transcription that did not overlap with annotated gene TSSs. Only the dREG coordinates that resided further than 1 kb from the annotated TSS were considered as transcribed enhancers. Transcription at enhancers was quantified by counting reads within dREG coordinates separately from plus and minus strands.

### Analysis of differential transcription of genes and enhancers

DESeq2 (Love *et al*., 2014) was used to perform differential expression analysis of genes and enhancers from PRO-seq data. Differential gene transcription was measured from gene bodies (+500 bp from TSS to -500 bp from CPS), whereas differential enhancer transcription was measured using the entire length of an enhancer from plus and minus strands separately. Statistically significantly changed genes and enhancers were called using p-value threshold of 0.05 and fold change threshold of 1.25 for upregulation and 0.8 for downregulation.

### Identification of transcription start nucleotides (TSNs, +1nts) and pause coordinates

Single-nucleotide resolution of PRO-seq allows determination of the exact +1nt and the Pol II pause nucleotide. In PRO-seq reads, the 5’-end nucleotides enrich at the precise +1nt, whereas the 3’-ends report the locations engaged Pol II. To identify the +1 nt, bedgraph files containing 5’ end nucleotides of sequenced reads were intersected with the regions around the annotated TSSs (-500 bp from TSS to +500 bp from TSS). The single nucleotide displaying the highest coverage of PRO-seq 5’-end coordinates was retained as the position of the +1nt. To identify Pol II pause nucleotides, the same approach was used, except 3’ end nucleotides of sequencing reads were intersected with the regions around annotated TSS.

### Motif analysis of enhancers

To analyze enrichment of motifs within enhancers, enhancer summits were first determined from the two +1nt coordinates at the opposite strands. 600 bp windows around enhancer summits were selected to perform motif analysis using MEME-ChIP online tool (Machanick and Bailey, 2011).

### Gene clustering and heatmaps

Differentially expressed genes from 30 min PRO-seq and 60 min mRNA-seq were plotted using ComplexHeatmap (Gu et al. 2016) and clustered with R function hclust. Genes that were differentially expressed in both conditions were retained and a pseudocount of -5 and 5 for infinite values was applied. Clustered genes with regard to their differential expression pattern were analysed for gene ontology. Gene ontologies were obtained using the Panther Classification System (Thomas et al. 2021).

### eCluster identification

To identify clusters of enhancers based on genomic proximity, we iteratively scanned position sorted enhancer coordinates and formed initial groups when a minimum of five enhancers resided within a 12.5 kb distance. These initial enhancer groups were extended if additional enhancers resided within a 2 kb distance from the preliminary group. At a subset of sites, which showed high enhancer occurrence, the iterative scanning produced overlapping groups of enhancers. These overlapping enhancer groups were combined using bedtools merge with -d 0 (Quinlan *et al*., 2010), producing 480 eClusters with a minimum of 5 eRPK Pol II density at both plus and minus strands. We further selected eClusters that contained at least five enhancers with a minimum eRPK 20 on both strands, yielding the final list of 404 eClusters (Supplementary Dataset 3). The framework for eCluster identification is provided in GitHub (https://github.com/Vihervaara/eClusterer).

### Transcriptional responses of clustered enhancers

To analyse responses of enhancers within eClusters, we compared the heat-induced change at individual enhancers (log2FC eRPK) to the mean change in the eCluster [mean(log2FC eRPKs in the eCluster)]. The fraction of enhancers that showed the same direction (log2FC > 0 | log2FC < 0) than the eCluster’s mean log2FC was recorded. The enhancer responses within eClusters were compared to two control groups that were formed from the unclustered enhancers: 1) position matched unclustered enhancers, and 2) transcription matched unclustered enhancers. The position matched enhancers resided adjacent to each eCluster. For each eCluster, we selected the same number of unclustered enhancers closest to the end of the eCluster (toward the end of the chromosome). For eClusters that resided close to a chromosome end, the position matched enhancers were selected from the start of the eCluster toward the chromosome start. For the expression matched group, we used the Sample package in R (https://rsample.tidymodels.org), randomly selecting unclustered enhancers from eRPK quantile bins, generating a list of unclustered enhancers with a matching eRPK distribution to clustered enhancers. These transcription matched enhancers were sorted into groups, recapitulating the total count of groups and individual enhancers within them as in the eClusters.

### eCluster sequence analyses

Fasta files were generated from enhancer coordinates using bedtools getfasta (Quinlan and Hall, 2010). The fasta files were analysed with Mash sketch (Ondov *et al*., 2016), using its smallest recommended kmer unit to identify all 9-mer sequences in each enhancer. Matching 9-mers were searched pairwise within clustered enhancers, and the counts of shared 9-mers and total 9-mers recorded. From the fraction of shared per total kmers, Mash uses MinHash and jaccard index approximation to compute a distance score with a value between 0-1 (Ondov *et al*., 2016). We converted the distance score to a measure of similarity (1 - distance), and generated correlation matrices to visualize the sequence similarity of enhancers within a cluster.

#### Genome files and annotations

CanFam6 annotation. Genes were categorized according to RefSeq annotation (https://www.ncbi.nlm.nih.gov/books/NBK21091/table/ch18.T.refseq_accession_numbers_an <u>d_mole/?report=objectonly</u>) to four categories, mRNA, predicted mRNA, ncRNA, and predicted ncRNA.

## Acknowledgements

We thank the members of the Vihervaara laboratory for valuable discussions. Juka and Juriaan Moonen are gratefully acknowledged for providing the photo in Figure 1A. This work was financially supported by Science for Life Laboratory (A.V., SciLifeLab Fellowship), Swedish Research Council (A.V., 2021-02668), and Royal Institute of Technology (A.V.).

## Author contributions

S.V.H. and A.V. conceived and designed the study. S.V.H. conducted the laboratory work. S.V.H., A.R., S.A., J.T. and A.V. conducted the computational analyses. All authors interpreted the results. S.H. and A.V. wrote the manuscript with help from all the authors.

---

**Supplementary figure 1.**
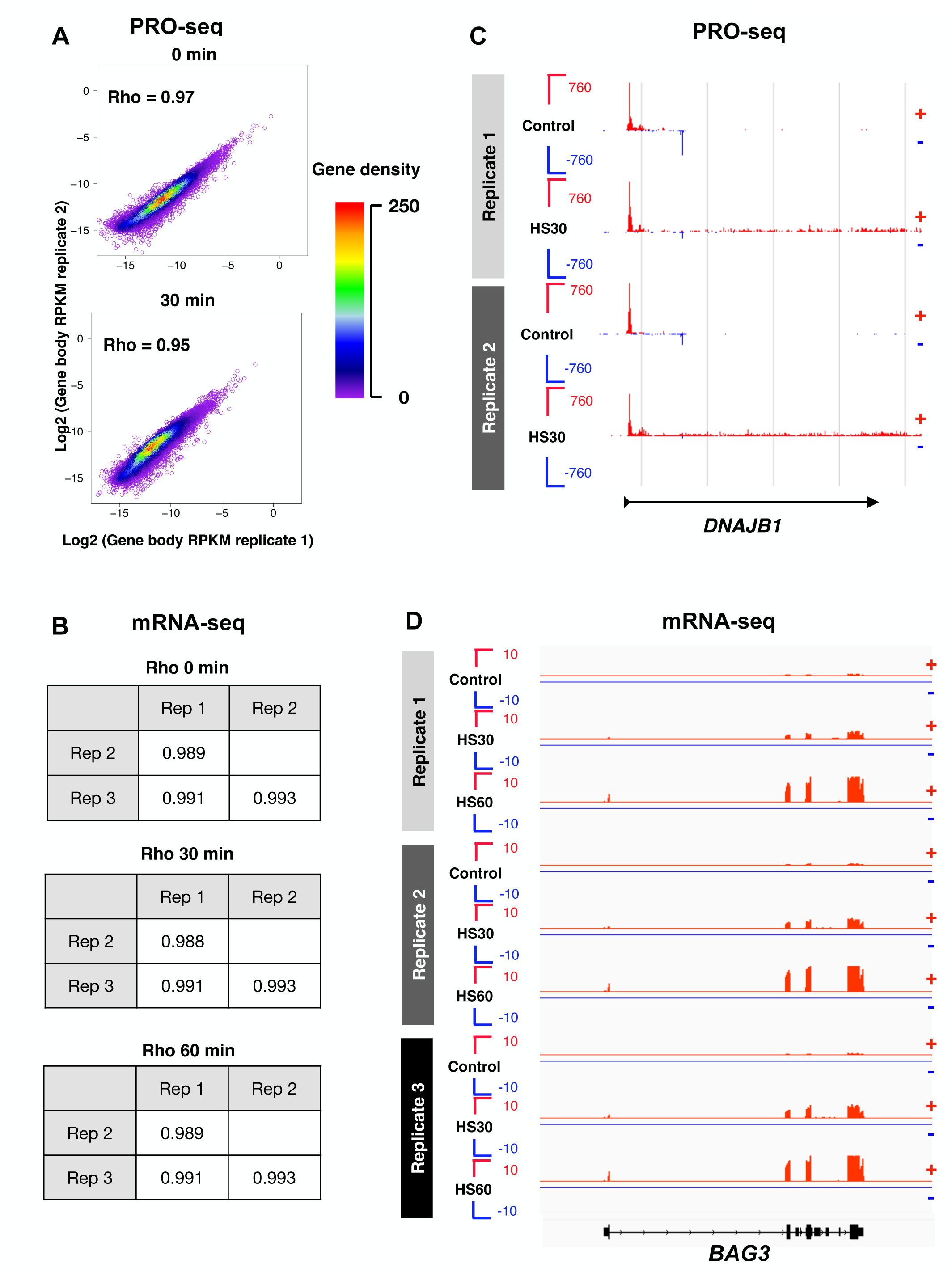
Correlations between PRO-seq and mRNA-seq replicates. **A)** Correlations between PRO-seq replicates. Genebody FPKM (reads per kilobase per million mapped reads) counts were correlated between biological replicates of control (0 min) and heat shock sample (30 min) to determine Spearman’s rank correlation coefficient (Rho). **B)** Correlations between mRNA-seq replicates. Correlations between biological replicates are determined from exonic read counts of mRNA-coding genes. **C-D)** Genome browser views of PRO-seq (C) and mRNA-seq (D) replicates across example genes.

**Supplementary figure 2.**
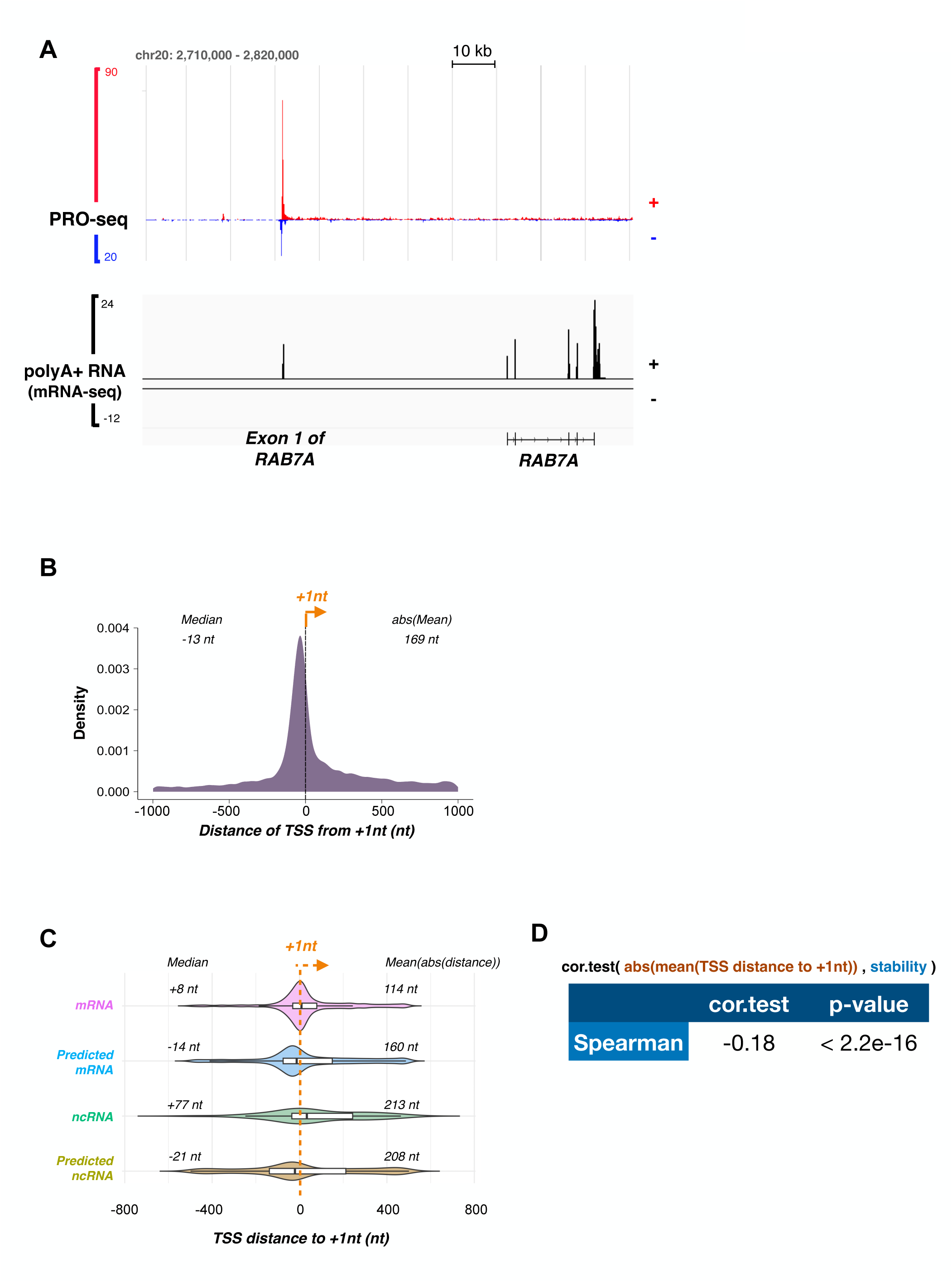
Distance of annotated TSS from the precise +1nt. **A)** Genome browser tracks of polyA+ RNA-seq and PRO-seq at *RAB7A* gene. The profile of transcription (PRO-seq) is shown to initiate tens of kbs upstream the annotated TSS. The presence of mRNA-seq signal at the transcription initiation supports the presence of an unannotated exon of *RAB7A*. **B)** Distance of annotated TSS from the +1nt across all dog genes. **C)** The distance between annotated TSS and the +1nt for distinct categories of genes. **D)** Spearman correlation measured between RNA stability (mRNA-seq FPKM / PRO-seq FPKM) and the TSS distance to the +1nt.

**Supplementary figure 3.**
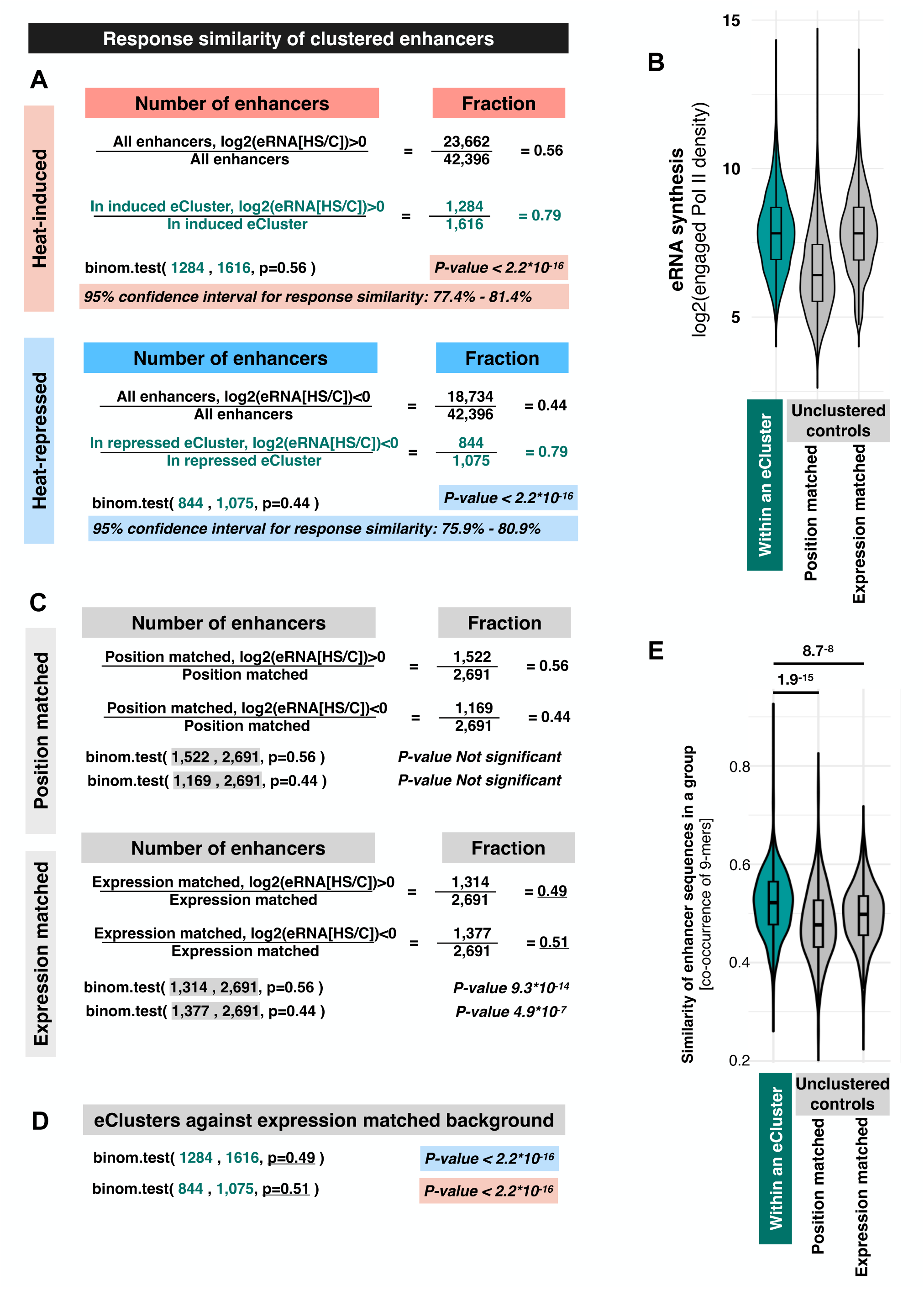
Individual enhancers within eClusters show unified responses. **A)** Fraction of enhancers showing induced (red) or reduced (blue) transcription. The binomial test of fraction enhancers in eClusters is compared to fraction similarly changed enhancers across the genome. The exact binomial calculations, p-values, and confidence intervals are shown. **B)** Enhancer transcription (eRPK) compared between the clustered enhancers (eClusters), and matched unclustered enhancers (position matched and expression matched unclustered enhancers). **C)** Fraction of unclustered enhancers and their binomial test conducted as in B. **D)** Binomial test of enhancer responses within eClusters calculated against the background obtained from the position matched or expression matched enhancers in C. **E)** Similarities of 9-mers with enhancers in eClusters (teal) or position matched (grey) or expression matched (grey) unclustered enhancers.

**Supplementary figure 4.**
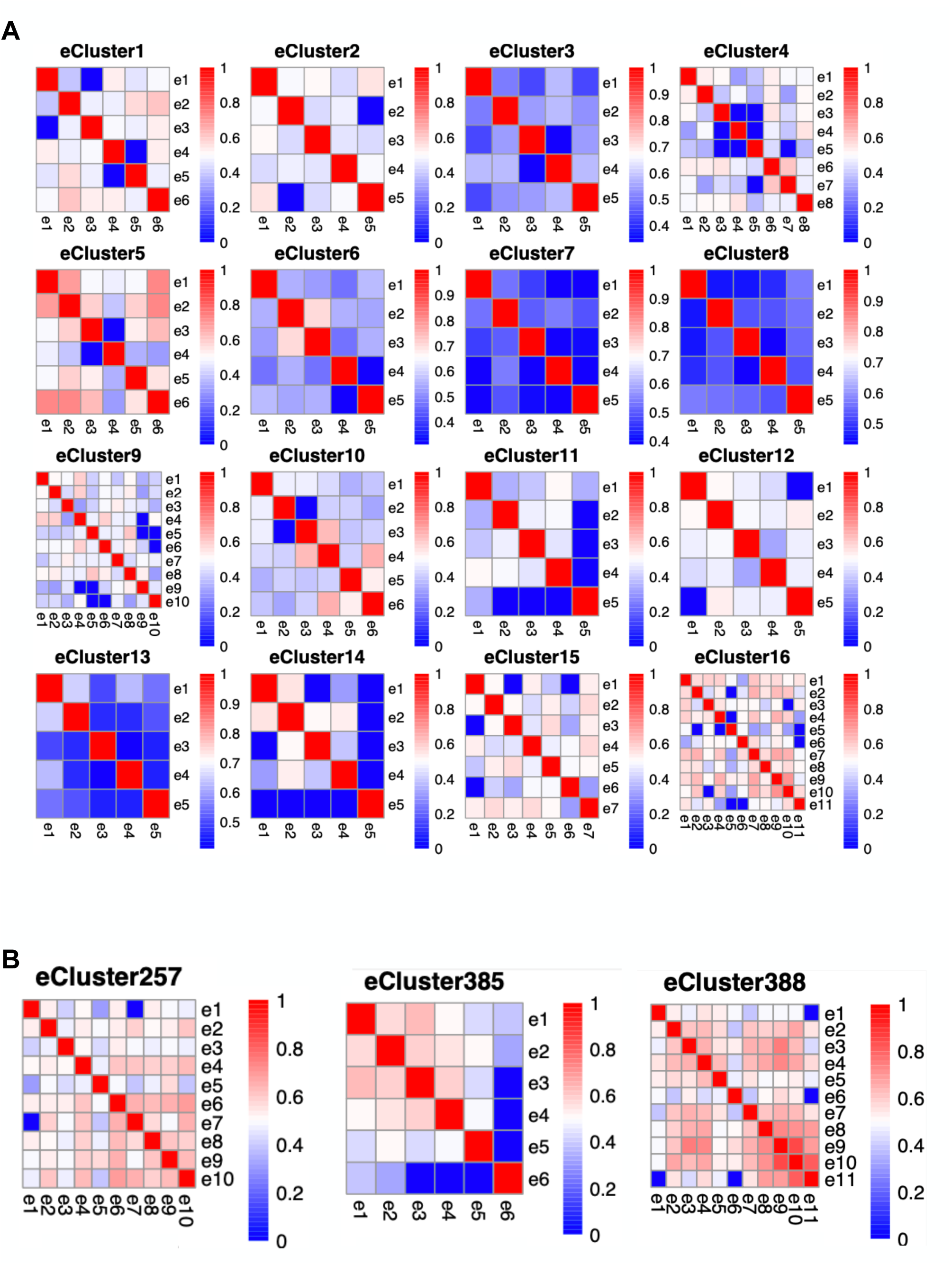
Unified response of enhancers within eClusters does not require sequence similarity. **A-B)** Pairwise comparison of 9-mers between enhancers within eClusters. The first 16 eClusters (by genomic position) are shown as examples (A), and compartmentalization of enhancer similarity is illustrated in selected eClusters (B).

